# Transcriptomic Analysis Identifies Transient Mesendodermal State and Lineage Divergence in Human Pluripotent Stem Cell Differentiation

**DOI:** 10.64898/2026.08.24.746642

**Authors:** Alice C. Borges, Mariana A. Branco, João P. Cotovio, Ana Rita Gomes, Joana E. Saraiva, Leonilde M. Moreira, Joaquim M.S. Cabral, Domingos Henrique, Maria Margarida Diogo, Tiago G. Fernandes

**Affiliations:** iBB - Institute for Bioengineering and Biosciences and Department of Bioengineering, Instituto Superior Técnico, Universidade de Lisboa, Lisbon, Portugal; Associate Laboratory i4HB Institute for Health and Bioeconomy, Instituto Superior Técnico, Universidade de Lisboa, Lisbon, Portugal; Gulbenkian Institute for Molecular Medicine (GIMM), Lisbon, Portugal; Collaborative Laboratory AccelBio, Biocant Park, Parque Tecnológico de Cantanhede, Cantanhede 3060-197, Portugal

**Keywords:** Comparative Transcriptomics, Human Pluripotent Stem Cells, Germ Layer Specification, Primitive Streak, Lineage Commitment

## Abstract

Human pluripotent stem cells serve as a vital model for studying early human lineage specification, yet conventional assessments relying on endpoint canonical markers of the three germ layers may overlook transient intermediate states and broader cellular programs. Here we combined directed differentiation of human induced pluripotent stem cells toward neuroectodermal, cardiac mesodermal, and hepatic endodermal lineages with comparative transcriptomic profiling across timepoints. Our analyses revealed a transient primitive streak-like mesendodermal state shared by mesodermal and endodermal trajectories, followed by lineage-specific divergence characterized by distinct transcriptional, metabolic, proliferative, and chromatin remodeling dynamics. Notably, endodermal differentiation exhibited rapid definitive endoderm commitment with enriched oxidative metabolism, whereas cardiac mesoderm differentiation showed progressive transcriptional remodeling and cardiac progenitor activation. These findings demonstrate that comparative transcriptomics can resolve developmental intermediates and cellular-state dynamics during human germ layer specification, providing a framework for evaluating lineage commitment beyond endpoint canonical marker expression, and to inform strategies for optimizing or redirecting differentiation.

## INTRODUCTION

Human embryogenesis is a tightly coordinated developmental process involving progressive changes in transcriptional programs, cellular identity, and tissue organization^1^. During early development, pluripotent epiblast cells, a population of cells in the early embryo, give rise to the three embryonic germ layers: ectoderm, mesoderm, and endoderm^2^. This process is initiated during gastrulation, when the primitive streak emerges and epiblast cells undergo lineage specification according to their position, signaling environment, and timing of ingression^3^.

In mammals, cells that move (*i.e.*, ingress) through the primitive streak contribute to mesodermal and endodermal lineages via a transient mesendodermal state^4^, characterized by the expression of developmental regulators such as T/Brachyury and Eomesodermin (EOMES)^5,6^. In contrast, epiblast cells that do not pass through the primitive streak primarily give rise to ectodermal lineages^7^. While many aspects of early development are conserved across species^8–11^, certain features of human embryogenesis are unique, underscoring the need for experimental models that can accurately capture early human lineage specification^1,11,12^.

Advances in stem cell biology and culture systems have greatly expanded the ability to study human development in vitro^13^. In particular, human pluripotent stem cells, including human induced pluripotent stem cells (hiPSCs), provide a tractable platform for modeling lineage commitment and organ development using both two-dimensional (2D) and three-dimensional (3D) differentiation systems. While 3D models better capture aspects of tissue organization and morphogenesis, 2D models offer controlled, reproducible, and temporally accessible systems to compare transcriptional changes across defined stages of differentiation. These approaches allow researchers to investigate developmental processes without the ethical and practical challenges associated with using human embryos directly^13–16^. Additionally, the increasing availability of large-scale gene expression (*i.e.*, transcriptomic) datasets has enabled comprehensive characterization of gene expression programs associated with pluripotency, differentiation, and lineage specification^17^.

Despite these advances, interpreting lineage commitment within complex differentiation trajectories remains challenging. Differentiation efficiency is commonly assessed using canonical lineage markers, a practice that simplifies the dynamic and continuous nature of development into discrete lineage identities. While this approach is useful for confirming target cell identity, it may overlook transient intermediate states and broader cellular programs that accompany lineage establishment. Transient states of cellular competence are of particular interest since they represent developmental windows in which cells retain plasticity, activate shared programs, and become progressively primed towards distinct lineage outcomes, essential to recapitulate and redirect developmental trajectories. Increasing evidence suggests that lineage specification is governed not only by the activation of lineage-specific transcription factors, but also by coordinated changes in broader cellular-state programs, such as metabolism, proliferation, chromatin remodeling, and signaling activity^18–22^. However, the extent to which directed differentiation protocols reproduce shared early developmental transcriptional programs, and how these programs diverge during specification toward distinct germ-layer lineages, remains incompletely understood.

To address these challenges, we combined directed differentiation of human pluripotent stem cells toward derivatives of the three embryonic germ layers, neuroectoderm, cardiac mesoderm, and hepatic endoderm, with comparative transcriptomic analysis across differentiation timepoints. Notably, mesoderm and endoderm both arise through a shared primitive streak-associated trajectory, making the resolution of this early mesendodermal state into distinct lineages of particular interest. By integrating temporal transcriptomic profiling with module-based analysis of developmental and cellular-state programs, we identified a transient primitive streak-like mesendodermal state shared by mesodermal and endodermal differentiation trajectories. Our analysis further suggests that lineage divergence is associated with distinct proliferative, metabolic, signaling, and chromatin-associated transcriptional states, with rapid endoderm commitment contrasting with more progressive cardiac mesoderm maturation. These findings highlight the value of analyzing differentiation trajectories beyond canonical lineage markers and support the use of comparative transcriptomics for assessing human lineage commitment in vitro.

## MATERIALS & METHODS

### Human cell lines

The human induced pluripotent stem cell (hiPSC) lines DF6-9-T.B, obtained from WiCell Bank, Wisconsin, USA, and F002.1A.13, obtained from TClab – Tecnologias Celulares para Aplicação Médica, Unipessoal, Lda, were used in this study. Cells were maintained under feeder-free conditions in humidified incubators at 37 °C and 5% CO₂. For routine hiPSC culture, cells were cultured in mTeSR™1 medium (STEMCELL Technologies™) supplemented 1:200 (v/v) with penicillin/streptomycin (Gibco™). Cells were plated on six-well plates coated with Matrigel^®^ Growth Factor Reduced Matrix (Corning^®^), and the medium was changed daily. Cells were passaged every 3–4 days, when they reached approximately 70% confluency, using 0.5 mM EDTA (Invitrogen™).

### Neural ectoderm differentiation

Neural induction was initiated when hiPSCs reached 90–100% confluency. The basal differentiation medium, N2B27, consisted of 50% (v/v) DMEM/F12/N2 and 50% Neurobasal/B27. DMEM/F12/N2 was prepared using DMEM/F12 medium (Thermo Fisher Scientific) supplemented with 1% (v/v) N2 supplement (Thermo Fisher Scientific), 1.6 g/L glucose (Sigma), 1% (v/v) penicillin/streptomycin, and 20 μg/mL insulin (Sigma). Neurobasal/B27 was prepared using Neurobasal medium (Thermo Fisher Scientific) supplemented with 2% (v/v) B27 supplement (Thermo Fisher Scientific), 2 mM L-glutamine (Thermo Fisher Scientific), and 1% (v/v) penicillin/streptomycin. Cells were cultured for 12 days in differentiation medium supplemented with 10 μM SB431542 and 100 nM LDN193189 (StemMACS™), with daily medium change.

### Cardiac-mesoderm differentiation

hiPSCs were differentiated into cardiac mesoderm according to the previously published protocol by Branco et al. (2019)^23,24^, with minor modifications if applicable. Briefly, for 2D monolayer culture, differentiation was initiated when cells reached 90– 95% confluency. RPMI 1640 medium (Gibco™) was used as the basal medium. On day 0, cells were cultured in basal medium supplemented with 2% (v/v) B-27 minus insulin (Gibco™) and 6 μM CHIR99021, a GSK3 inhibitor (Stemgent). After 24 h, the medium was changed to RPMI 1640 supplemented with 2% B-27 minus insulin. From day 3 to day 5, cells were cultured in basal medium supplemented with 5 μM IWP-4, a Wnt inhibitor (Stemgent). From day 7 onward, cells were cultured in RPMI 1640 supplemented with 2% (v/v) B-27 (Gibco™).

### Hepatic endoderm differentiation

For differentiation into hepatic endoderm, RPMI 1640 medium (Gibco™) was used as the basal medium. RPMI 1640 was supplemented with 1% (v/v) B-27™ minus insulin (Gibco™) and 1:200 (v/v) penicillin/streptomycin (Gibco™). hiPSCs were previously expanded and seeded in twelve-well plates coated with iMatrix-511 (Nippi/Matrixome^®^) at a density of 4 × 10⁵ cells/well. Cells were maintained in mTeSR™1 supplemented 1:200 with penicillin/streptomycin, and the medium was changed daily until cells reached 90–95% confluency.

To initiate definitive endoderm differentiation, basal medium was supplemented with 100 ng/mL Activin A (PeproTech), 1 mM sodium butyrate (NaB; Sigma-Aldrich^®^), and 2 μM CHIR99021 (CHIR; Stemgent™). On days 1 and 2, cells were cultured in basal medium supplemented with 100 ng/mL Activin A. On day 3, to induce hepatic specification, medium was additionally supplemented with 10 ng/mL FGF2 (PeproTech) and 20 ng/mL BMP4 (PeproTech). On day 6, cells were transferred to hepatocyte culture medium (HCM) BulletKit™ medium (Lonza), without human epidermal growth factor (hEGF), supplemented with 10 ng/mL oncostatin M (OSM; R&D Systems™), 0.1 μM dexamethasone (Dex; Sigma-Aldrich^®^), and 20 ng/mL hepatocyte growth factor (HGF; Sigma-Aldrich^®^).

### RNA sequencing Analysis

#### RNA extraction

Total RNA was extracted from cells at different stages across the three germ layer differentiation protocols using the High Pure RNA Isolation Kit (Roche), according to the manufacturer’s instructions.

#### Bulk-RNA sequencing and data processing

Libraries were prepared using the Lexogen QuantSeq 3′ mRNA-Seq Library Prep Kit FWD for Illumina, according to the manufacturer’s standard protocol. Sequencing was performed on Illumina HiSeq or NextSeq platforms, using 50-cycle or 75-cycle protocols, respectively. Three biological replicate samples (n=3) were analyzed for each differentiation timepoint.

For endoderm and mesoderm samples, read quality assessment, read mapping, and gene counting were performed using the standard BlueBee Genomics Platform pipeline. For each sample, the pipeline included read trimming with BBDuk, alignment to the reference genome using STAR, gene-level read counting with HTSeq, and quality control using RSeQC.

For ectoderm samples, raw RNA-sequencing reads in FASTQ format were processed using the R/Bioconductor environment (version 4.4.2). Reference genome indexing and read alignment were performed using the Rsubread package (version 2.20.0)^25^ against the human GRCh38 primary assembly obtained from Ensembl release 115. Gene-level read quantification was generated from aligned reads using featureCounts with Ensembl GRCh38 gene annotations. Gene identifiers were converted from Ensembl IDs to HGNC gene symbols using the biomaRt package and the Ensembl hsapiens_gene_ensembl dataset. Genes without corresponding HGNC annotations were excluded from downstream analyses. For genes mapping to multiple Ensembl identifiers, read counts were collapsed by summing counts across shared HGNC symbols.

Quality control analyses were performed to assess sequencing complexity and sample distribution. Cumulative read distribution plots and density plots of log10-transformed read counts were generated for all samples and experimental groups. Lowly expressed genes were filtered before normalization based on total read abundance across samples.

#### Differential Expression Analysis

Normalization and differential expression analyses were performed using the edgeR and limma R packages. Raw count matrices were converted into DGEList objects and normalized using the trimmed mean of M-values method. Mean–variance relationships were modeled using the voom transformation, generating log2-transformed counts per million values and precision weights for linear modeling. Differential gene expression between experimental timepoints was assessed using linear modeling and empirical Bayes moderation implemented in limma. Contrasts were generated relative to the D0 condition for each developmental stage analyzed. Differentially expressed genes were ranked according to log2 fold change and statistical significance.

Gene Ontology (GO) enrichment analysis and Gene Set Enrichment Analysis (GSEA) were performed using the clusterProfiler R package. For Gene Ontology analysis, the p-value threshold was set to 0.05. For Gene Set Enrichment Analysis, a p-value cut-off of 0.05 was used, and p-values were adjusted using the Benjamini– Hochberg method.

#### Dimensionality Reduction Analysis

Dimensionality reduction analyses were performed using an integrated transcription factor expression matrix generated from ectoderm, mesoderm, and endoderm RNA-sequencing datasets. After preprocessing and normalization, genes shared across all three germ layer datasets were identified and merged into a combined expression matrix. Transcription factor-associated genes were retrieved from the C3 collection of Molecular Signatures Database (MSigDB)^26^ using the msigdbr R package (version 26.1.0), and the integrated matrix was subset to retain only these genes.

Normalized expression values were generated using the voom transformation implemented in limma and averaged across biological replicates for each developmental condition. The resulting expression matrix was transposed so that samples represented observations and genes represented variables. Expression values were then standardized using z-score scaling before dimensionality reduction.

PCA was performed in R using the prcomp function with variance scaling enabled. Principal component coordinates were extracted and annotated by germ layer lineage and developmental stage for visualization.

UMAP was performed using the uwot R package on the scaled transcription factor expression matrix, with n_neighbors set to 5 and min_dist set to 0.3. To ensure reproducibility, a fixed random seed was applied using set.seed (= 42).

PCA and UMAP plots were generated using ggplot2. Samples were colored according to lineage classification, and developmental stage labels were added where appropriate.

#### Module Scoring Analysis

Module scores were calculated for each sample as the average z-scored expression of genes within each module. Scores were computed across developmental timepoints and lineages using normalized expression matrices derived from voom-transformed RNA-sequencing data. To assess lineage-associated cellular states independently of the PC2-derived module, gene sets representing glycolysis, oxidative phosphorylation (OXPHOS), cell cycle progression, stemness, and chromatin remodeling were derived from significantly enriched GO biological process terms identified among genes with negative PC2 loading in the endoderm-mesoderm comparison. The stemness module was defined using the published Ben-Porath embryonic stem cell gene set^27^. Complete gene lists for all modules are provided in Supplementary Table 1.

For each module, expression values were standardized by gene-wise z-score transformation before score calculation. Module scores were subsequently compared across lineages and developmental stages. Module scores were analyzed using linear models (lm) (equivalent to a factorial ANOVA) with lineage, differentiation day, and their interaction as fixed effects (Score ∼ Lineage × Day). Pairwise comparisons between lineages within each differentiation day were obtained using the emmeans package (version 1.1.12) to estimate marginal means. P values < 0.05 were considered statistically significant. Data visualization was performed in R using ggplot2.

#### Developmental Gene Trajectory Analysis

Genes contributing most strongly to lineage separation were identified from the positive and negative loadings of PC2 obtained from PCA performed on samples from D3 onwards. Normalized expression values derived from the voom-transformed RNA-sequencing matrix were extracted for these gene sets and organized according to developmental day and lineage identity. For each lineage and developmental stage, the expression module score across genes within each PC2-associated gene set was calculated and used to generate temporal trajectory plots.

For visualization, individual genes were plotted as line traces connecting sequential developmental stages within the mesoderm and endoderm lineages. Replicate samples were retained to preserve temporal variability within each condition, and gene trajectories were faceted by lineage.

Dynamics visualization and data handling were performed in R using ggplot2 and associated tidyverse packages.

### Accession numbers

RNA-seq data for this study are available through Gene Expression Omnibus (GEO) Accession number GSE116574 (Mesoderm), GSE311252 (Endoderm) and GSE338003 (Ectoderm).

## RESULTS

### Directed differentiation of hiPSCs into mesoderm and ectoderm captures a transient primitive streak-like mesendodermal state

Transcriptomic data from in-house 2D hiPSC differentiation experiments toward ectodermal, mesodermal and endodermal lineages were curated against previously published differentiation protocols^23,28–30^ (Fig. 1A) to retain lineage-specific genes while minimizing protocol-dependent signatures. The resulting curated gene set was used for dimensionality reduction analyses.

**Figure 1A.**
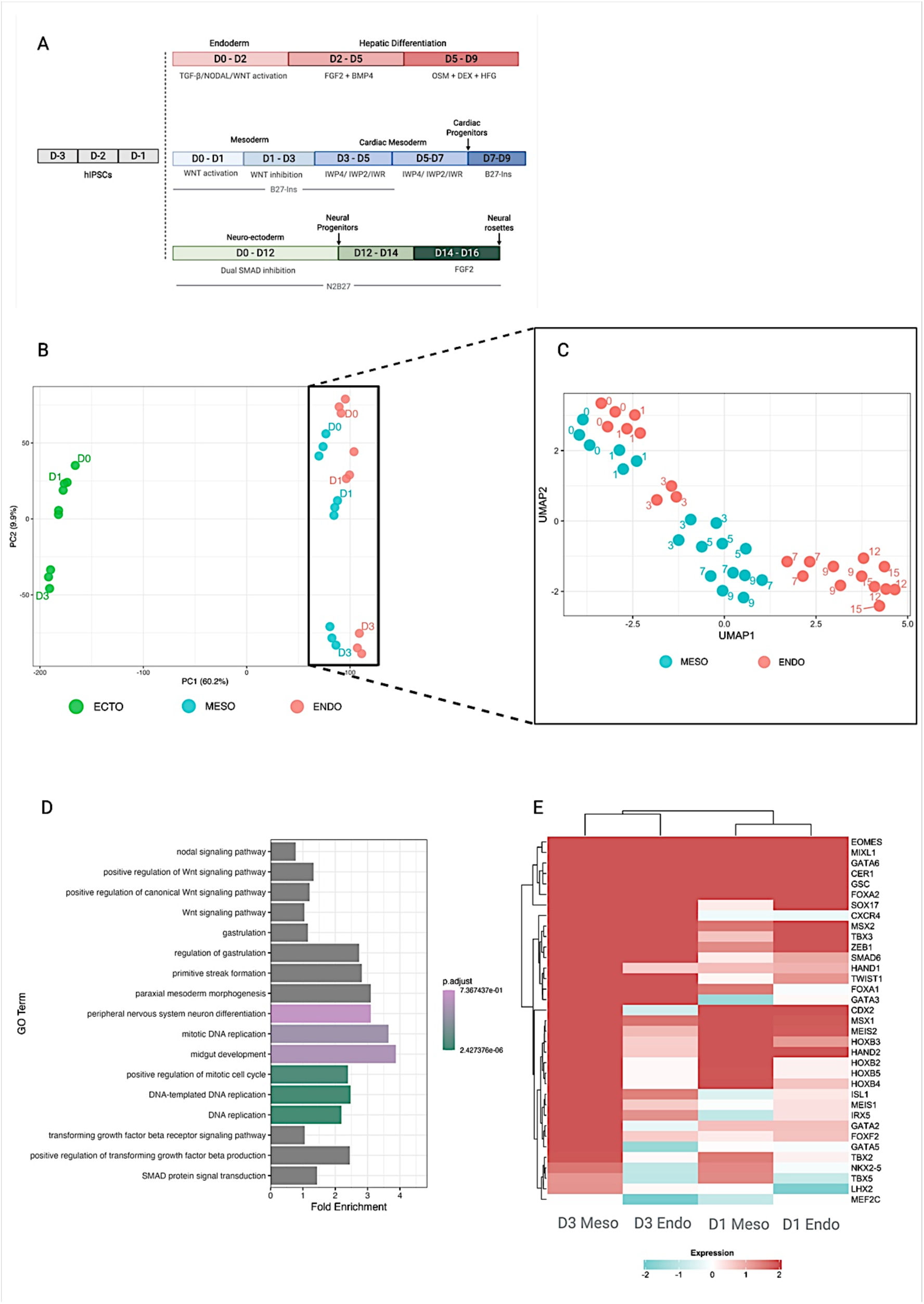
Schematic of the directed 2D differentiation protocols used to generate endodermal, mesodermal/cardiac, and neuroectodermal lineages from hiPSCs: Colored bars denote differentiation stages and sampling intervals; key signaling factors are indicated below each stage. **Figure 1B | PCA visualization of lineage trajectories during 2D hiPSC differentiation toward ectoderm, mesoderm and endoderm:** Samples are colored by lineage (mesoderm, green; endoderm, blue; ectoderm, red) and labeled by differentiation day. PC1 and PC2 explain 60.2% and 9.9% of variance, respectively. **Figure 1C | UMAP visualization of lineage trajectories during 2D hiPSC differentiation toward mesoderm and endoderm:** UMAP projection of transcriptomic profiles from in-house 2D hiPSC differentiation toward mesoderm (Meso, Blue) and endoderm (Endo, red). Points represent samples, colored by lineage, with numbers indicating differentiation day. Early time points from both lineages cluster together, followed by divergence along distinct mesodermal and endodermal trajectories. **Figure 1D | GO enrichment of PC2 least contributing genes highlights cell cycle, gastrulation and embryonic patterning programs:** Bars show fold enrichment and are colored by adjusted P value. **Figure 1E | Lineage marker expression during early differentiation:** Heatmap of logFC values for curated mesodermal, endodermal, and primitive streak marker genes at Day 1 and Day 3 of differentiation. Rows represent genes and columns represent lineage-specific samples. Colors denote relative expression levels.

PCA of the three differentiation trajectories revealed clear separation of ectodermal samples from mesodermal and endodermal samples (Fig. 1B, Supp Fig. 1A). Genes contributing to PC1 were strongly enriched for developmental commitment programs, with positive loadings associated with mesodermal development and negative loadings enriched for neurodevelopmental and brain-associated processes (Supp Fig. 1B, 1C). In contrast, PC2 primarily captured temporal progression across differentiation, with negative loadings corresponding to earlier developmental stages and showing significant enrichment for embryonic organ development pathways (Supp Fig. 1D).

The marked segregation of ectodermal samples from mesodermal and endodermal samples is consistent with gastrulation-associated lineage relationships, in which ectodermal lineages arise from epiblast cells that do not ingress through the primitive streak. In contrast, the closer association between mesodermal and endodermal samples suggests the presence of a shared transcriptional program consistent with their common mesendodermal origin^4,5,7^. To further investigate this relationship, we applied Uniform Manifold Approximation and Projection (UMAP), a dimensionality reduction technique that preserves local and global data structure, focusing specifically on mesodermal and endodermal lineages. UMAP visualization revealed that samples from both lineages clustered together at early differentiation timepoints (Fig. 1C), supporting a transient shared transcriptional state during early lineage specification. From day 3 (D3) onward, samples began to segregate according to lineage, suggesting the emergence of lineage-specific transcriptional divergence.

This divergence was further supported by PCA, which showed progressive separation of mesodermal and endodermal samples during differentiation (Supp Fig. 1E). PC1 primarily captured differentiation progression, whereas PC2 reflected lineage divergence. Genes with high positive loadings on PC2 were significantly enriched for mesodermal and cardiac developmental processes, including heart morphogenesis, cardiac muscle differentiation, chamber formation, mesenchymal development, and embryonic patterning (Supp Fig. 1F), indicating that PC2 captures the emergence of mesoderm-specific transcriptional programs. In contrast, genes with negative PC2 loadings were enriched for oxidative phosphorylation, translation, ribosomal function, and mitochondrial pathways, suggesting that endodermal samples are associated with a distinct metabolic and biosynthetic transcriptional state relative to mesodermal samples (Supp Fig. 1G).

Given the early clustering observed by UMAP and the lineage separation captured by PC2, we next investigated genes that did not contribute strongly to PC2-driven separation, reasoning that these genes may represent a shared transcriptional program between mesodermal and endodermal trajectories. PC2 loadings were extracted from the PCA, and genes were ranked by the absolute magnitude of their loadings. Those with absolute PC2 loadings within the lowest 10% of distribution (|loading| ≤ 10^th^ percentile) were selected as genes contributing least to lineage separation, resulting in a set of 1421 genes for downstream analysis. Enrichment analysis of this gene set revealed significant overrepresentation of pathways related to mitotic regulation, DNA replication, gastrulation, primitive streak formation, and pattern specification (Fig. 1D). These findings suggest that the differentiation protocols recapitulate early mesendodermal transcriptional programs characterized by high proliferative activity and features associated with gastrulation-like, pre-divergence stages^5,31–33^.

To further assess the shared mesendodermal identity, we examined the expression of known primitive streak, mesodermal, and endodermal markers using heatmap analysis of differentially expressed genes at day 1 (D1) and day 3 (D3) of differentiation (Fig. 1E). Consistent with the UMAP analysis, D1 samples from both lineages clustered together, with D3 endodermal samples remaining closer to the early D1 state, whereas D3 mesodermal samples showed greater divergence. The heatmap revealed a cluster of genes upregulated in both lineages at D1 that became downregulated in endodermal differentiation but remained sustained during early mesodermal differentiation (*CDX2, MSX1, MEIS2, HOXB3, HAND3*), supporting rapid divergence from a common early transcriptional state. This pattern is consistent with developmental dynamics in which primitive streak-associated programs, including transcriptional features linked to epithelial-to-mesenchymal transition and cell ingression in vivo, are transiently activated before mesodermal and endodermal lineage specification^7,33,34^.

In addition, a subset of transcription factors remained upregulated in both lineages during early differentiation (*EOMES, MIXL1, GATA6, CER1, GSC*), further supporting a shared primitive streak/mesendodermal origin (Fig. 1E). Tissue enrichment analysis reinforced these findings, showing enrichment for cardiac tissues, mesoderm, endoderm, and early embryonic structures, including blastocyst and trophoblast (Supp Fig. 1H). These results are consistent with the activation of an early developmental transcriptional program preceding lineage commitment.

Together, these results indicate that, despite the use of directed 2D differentiation protocols toward cardiac mesoderm and hepatic endoderm, both lineages transiently acquire a shared primitive streak/mesendodermal transcriptional state consistent with their common developmental origin in vivo.

### Transient metabolic and proliferative state divergence distinguishes early mesodermal and endodermal differentiation

Building on the separation observed at D3 by UMAP, we further examined mesodermal and endodermal divergence along PC2 to identify biological processes underlying this distinction (Fig. 1C - UMAP, Supp Fig. 1E – PCA). Genes with the strongest negative loadings on PC2, hereafter referred to as PC2-DOWN genes, were enriched for pathways associated with oxidative phosphorylation (OXPHOS), translation, ribosomal function, mitochondrial assembly, and mitochondrial metabolism (Supp Fig. 1G). This led us to ask whether endodermal differentiation is associated with a distinct metabolic or proliferative transcriptional state beyond canonical lineage-specification programs, which were enriched among genes with positive PC2 loadings. To quantify these patterns, we computed module scores using a curated gene set derived from the fifteen most enriched PC2-DOWN GO terms (Supp Fig. 1G), as well as independent metabolic and cell-cycle gene sets, across differentiation timepoints and lineages (Fig. 2A). At D1, endodermal samples showed a slightly greater reduction in PC2-DOWN module scores relative to baseline, suggesting an earlier but subtle shift away from pluripotency-associated metabolic and biosynthetic programs compared with mesodermal samples (Fig. 2B). This divergence was transient, as both lineages converged by D3 toward a similarly reduced metabolic/translation-associated state relative to pluripotent D0 cells (Fig. 2B). These results suggest that endodermal differentiation may undergo an earlier transition toward a less proliferative and more lineage-committed transcriptional state.

**Fig 2A.**
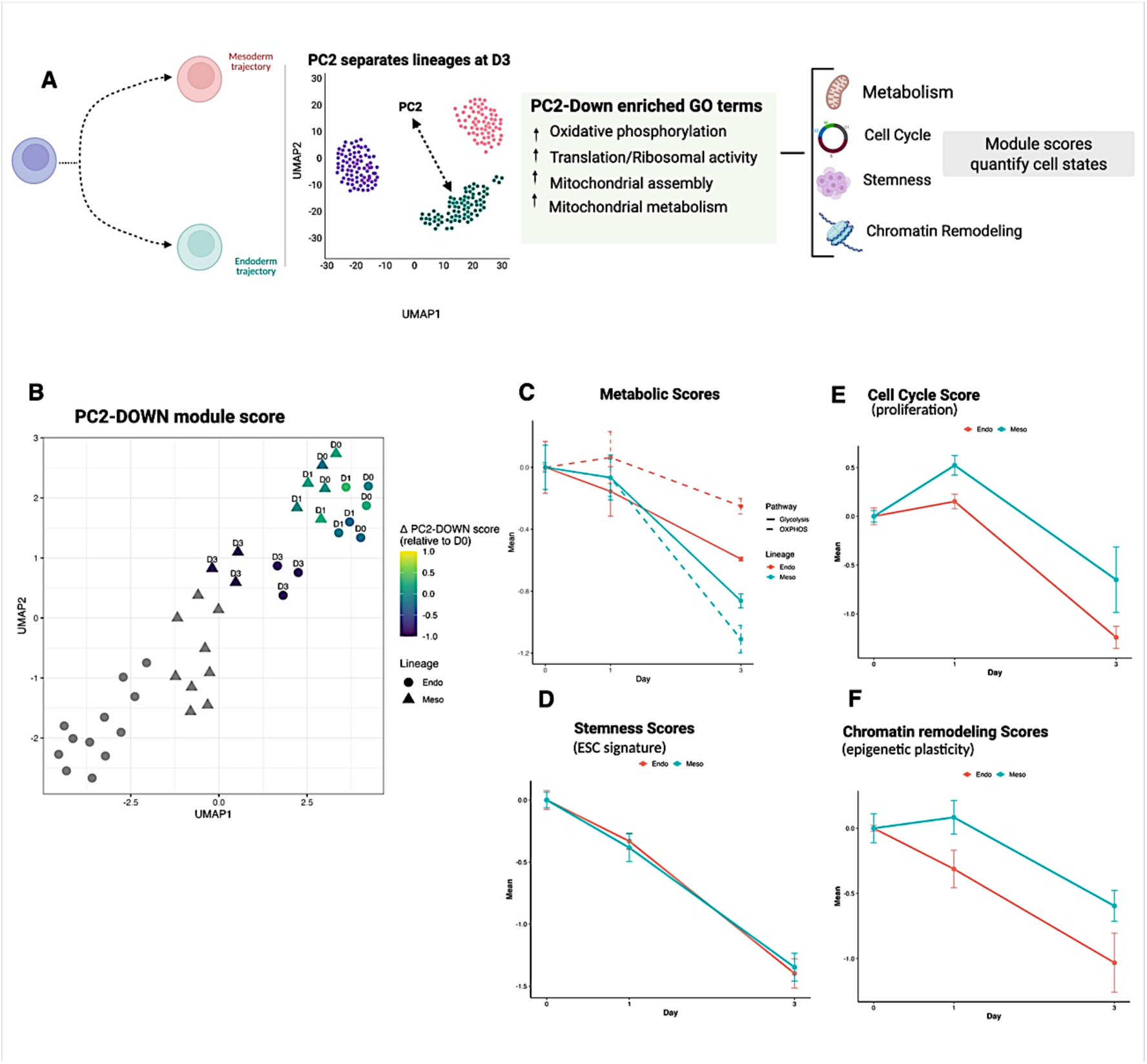
Schematic of the workflow used to derive biologically interpretable transcriptional modules from principal component loadings. **Fig 2B | UMAP representation of all samples colored by PC2-DOWN module score relative to the lineage-specific day 0 baseline.** Colors indicate module activity, shapes denote lineage, and labels indicate differentiation stage. **Fig 2C – F | (C–F) Temporal dynamics of glycolysis/OXPHOS (**C**), stemness (**D**), cell-cycle (**E**), and chromatin remodeling (**F**) module scores during differentiation.** Module scores were calculated as the mean z-score of genes within each signature and normalized relative to the lineage-specific day 0 baseline. Endoderm samples are shown in red and mesoderm samples in blue. Points indicate mean values and error bars represent SEM across biological replicates. Statistical comparisons between lineages at each time point were performed using a linear model followed by estimated marginal means (emmeans).

To further investigate the cellular processes contributing to this separation, we examined module scores associated with metabolism, cell-cycle progression, stemness, and chromatin remodeling. Metabolic and cell-cycle gene sets were derived from the enriched GO terms described above, and included genes associated with S-phase licensing (e.g., *MCM2*, *POLA1*, and *PCNA*), G1/S transition (e.g., *CCND1*, *CCNE1*, *CDK4*, *CDK6*, and *CDK2*), G2/M entry (e.g., *CDK1* and *CCNB1*), and checkpoint control (e.g., *CHEK1*, *RB1*, and *CDC25A*) (Supp Table 1). Stemness was assessed using a panel of embryonic stem cell-associated markers derived from the Ben-Porath gene sets^27^. Given the known association between glycolytic metabolism and pluripotency, and between oxidative phosphorylation and differentiation progression^21^, we also calculated glycolysis and OXPHOS module scores. The OXPHOS score was calculated using structural subunits of electron transport chain complexes I–V, excluding genes associated with mitochondrial biogenesis and ribosomal function.

In both lineages, glycolysis and OXPHOS-associated transcriptional scores decreased along the D0-to-D3 differentiation axis (Fig. 2C), consistent with a strong effect of differentiation time on both metabolic programs. Glycolytic scores declined similarly in mesodermal and endodermal cells, with no significant lineage-dependent differences. In contrast, OXPHOS-associated scores displayed significant effects of lineage and lineage-by-day interaction, reflecting distinct temporal trajectories between the two lineages. By D3, endodermal samples retained significantly higher OXPHOS scores than mesodermal samples (P < 0.0001), consistent with a relative shift toward oxidative metabolism during endoderm specification (Supp Fig. 1G). Together, GO enrichment and pathway scoring analyses suggest that endodermal differentiation is associated with a relative shift toward oxidative phosphorylation and electron transport chain-associated transcriptional programs compared with glycolytic metabolism.

Stemness-associated transcriptional scores decreased similarly in both lineages, with no significant differences between mesodermal and endodermal samples at any differentiation stage (all P > 0.69), indicating a comparable loss of pluripotency during lineage commitment (Fig. 2D). Analysis of cell-cycle-associated scores also revealed comparable proliferative states in mesodermal and endodermal samples at D0–D1, consistent with the proliferative nature of early primitive streak-associated differentiation stages (Fig. 2E)^35,36^. The decreasing proliferative state trend in endoderm at D1 did not reach statistical significance. However, by D3 endodermal samples showed a significant reduction in cell-cycle-associated transcriptional activity relative to mesodermal samples (P = 0.022), suggesting a more pronounced reduction in proliferative activity during endodermal differentiation. These findings suggest that endodermal cells progressively reduce cell-cycle-associated transcriptional activity during differentiation, with significant divergence from mesoderm becoming apparent by D3. This reduction occurs while endoderm maintains relatively higher oxidative metabolism-associated scores, consistent with a metabolic state linked to lineage patterning rather than solely cell division.

To further assess whether endodermal differentiation reaches a more stabilized lineage state by D3, we examined chromatin remodeling-associated transcriptional scores. Because chromatin remodeling is closely linked to cell-identity specification^6,22,37^, reduced chromatin remodeling scores may reflect decreased transcriptional plasticity following lineage commitment, although activation of specific differentiation programs may still require targeted chromatin remodeling. Endodermal samples displayed lower chromatin remodeling scores, with the difference reaching statistical significance at D3 compared with mesodermal samples (Fig. 2F, P = 0.047). This pattern is consistent with earlier stabilization of fate-associated chromatin programs in endoderm. In contrast, the relatively higher chromatin remodeling-associated scores in mesodermal samples suggest ongoing transcriptional and epigenetic remodeling during progressive cardiac mesoderm specification^22,37^.

Overall, these results suggest that the lineage separation observed by D3 reflects not only lineage-specific transcriptional programs, but also distinct cellular-state dynamics between more established endodermal cells and progressively specifying mesodermal cells.

### Distinct transcriptional trajectories reveal rapid definitive endoderm commitment and progressive cardiac mesoderm maturation

Having identified the transcriptional programs underlying lineage separation, we next examined the temporal behavior of the genes contributing most strongly to this divergence. Genes contributing to lineage separation displayed distinct temporal dynamics in each lineage. Endoderm-associated gene programs reached maximal activity by the lineage bifurcation point (D1) and changed comparatively little thereafter, whereas mesoderm-associated program continued to increase until D5 before stabilizing, indicating a more progressive transcriptional remodeling process (Fig. 3A). Therefore, we proceeded with the analysis of expression dynamics from individual transcription factors associated with each lineage-specific program.

**Fig 3A.**
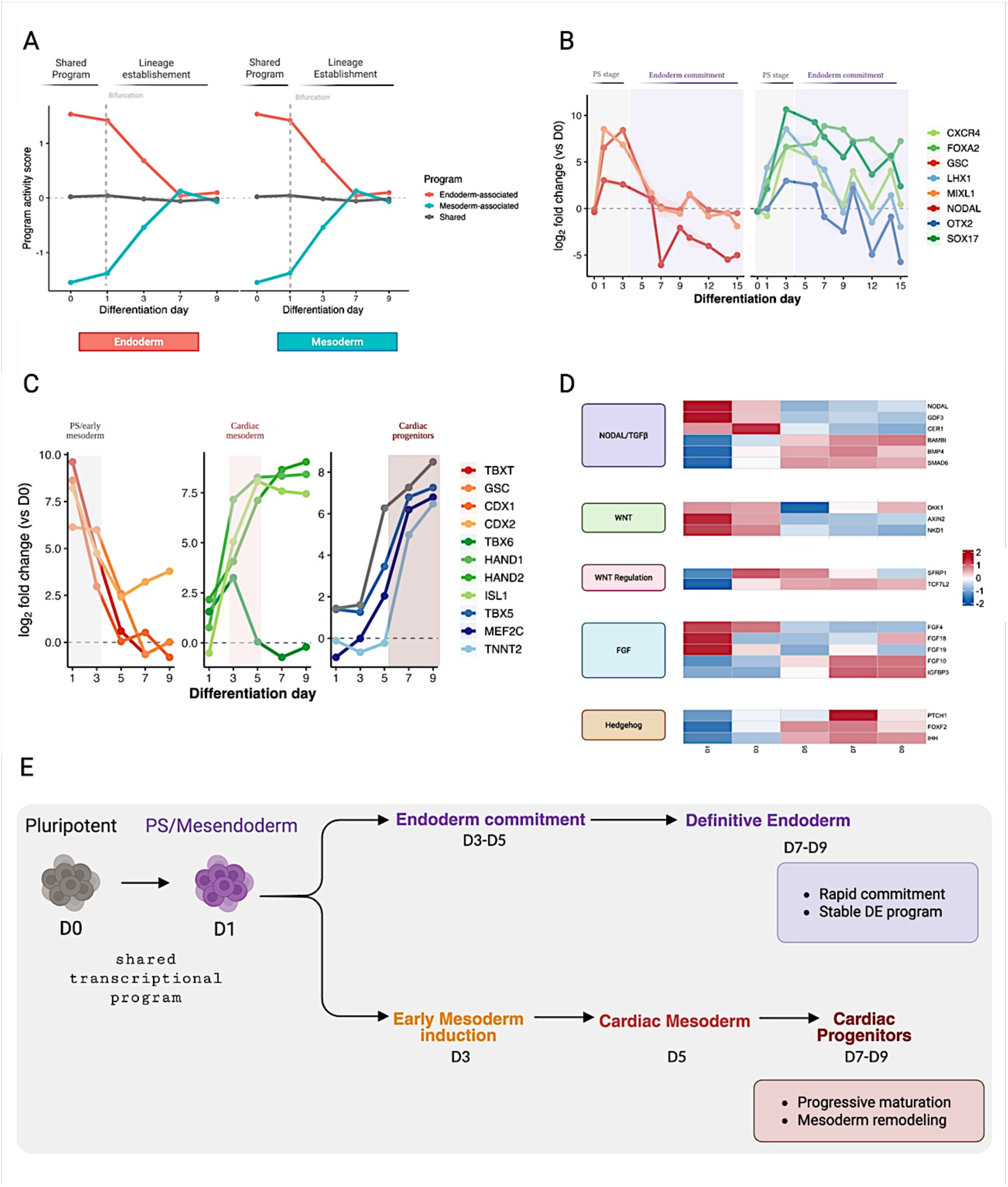
Temporal dynamics of PCA-derived shared, endoderm-associated, and mesoderm-associated transcriptional programs. Values represent median gene-wise standardized expression. The dashed line indicates lineage bifurcation. **Fig 3B | Endoderm lineage marker dynamics during differentiation.** Expression trajectories of representative primitive streak-associated (*NODAL, MIXL1, GSC*) and definitive endoderm-associated (*SOX17, FOXA2, CXCR4, LHX1, OTX2*) markers across the endoderm differentiation time course. Values represent mean log₂ fold-change relative to D0; shaded regions indicate ± SD across replicates. The transition from the shared primitive streak stage (D0–D3) to endoderm commitment is highlighted. **Fig 3C | Mesoderm and cardiac lineage marker dynamics during differentiation.** Temporal expression profiles of representative primitive streak/early mesoderm (*TBXT, GSC, CDX1, CDX2*), cardiac mesoderm (*TBX6, HAND1, HAND2, ISL1*), and cardiac progenitor (*TBX5, MEF2C*) and cardiomyocyte (*TNNT2)* markers during mesoderm differentiation. Values represent mean log₂ fold-change relative to D0. Shaded regions indicate the transition from primitive streak/early mesoderm to cardiac mesoderm and cardiac progenitor specification stages. **Fig 3D | Dynamic regulation of signaling pathways during mesoderm differentiation.** Heatmap showing gene-wise standardized expression (Z-scores) of representative components of the TGFβ/SMAD, WNT, FGF and Hedgehog pathways across differentiation. Genes are grouped by pathway and ordered according to temporal expression dynamics. Red indicates relative upregulation and blue indicates relative downregulation. **Fig 3E | Schematic summary of lineage progression during directed differentiation.** Pluripotent cells transition through a shared primitive streak/mesendoderm stage before diverging toward endoderm and mesoderm fates. Endoderm samples progress through definitive endoderm commitment, whereas mesoderm samples transition through cardiac mesoderm and cardiac progenitor stages. Developmental stages are shown according to established lineage markers and transcriptomic trajectories identified in this study.

In the endodermal trajectory, we tracked the expression of *SOX17*, *MIXL1*, and *GSC*, which are associated with primitive streak, mesendodermal, and endodermal specification programs (Fig. 3B). As expected, these markers peaked at D3, coinciding with the establishment of the endodermal lineage in our 2D differentiation system. *NODAL* and other members of the TGF-β superfamily also showed peak expression at D3. This is consistent with the established role of NODAL signaling in primitive streak formation and definitive endoderm specification, including the induction of key regulators such as *SOX17* and *FOXA2*^38,39^. In line with previous reports, we also observed a D3 expression peak for *CXCR4*, also known as CD184, which encodes a cell-surface chemokine receptor commonly used as a marker of definitive endoderm cells^40^ (Fig. 3B). Additional genes peaking at D3 included *LHX1*, also known as *LIM1*, and *OTX2* (Fig. 3B). These genes encode transcription factors associated with early embryonic patterning and endoderm-related developmental programs^41,42^. Interestingly, recent work has shown that *EOMES*, acting downstream of NODAL signaling, can activate *LHX1* in the epiblast, contributing to definitive endoderm lineage specification^43^.

To generate cardiomyocytes from mesodermal progenitors, differentiation was performed using a previously described protocol based on canonical WNT signaling activation from D0 to D1, followed by WNT inhibition from D3 to D5^23,24^. At early differentiation stages, gene set enrichment analysis revealed that D1-upregulated genes were significantly enriched for early developmental processes, including gastrulation, somitogenesis, pattern and axis specification, and mesoderm differentiation (Supp Fig. 2A). Consistent with our previous observation that endodermal and mesodermal trajectories share an early transcriptional program, canonical primitive streak and early mesoderm-associated markers, including *TBXT* and *GSC*, were strongly induced^44^. In addition, CDX transcription factors, including *CDX1* and *CDX2*, were upregulated, consistent with activation of posterior primitive streak-associated patterning programs during early mesoderm induction^45^ (Fig. 3C).

Early lateral plate mesoderm and cardiac mesoderm-associated transcriptional programs became evident around day 3, marked by the upregulation of regulatory transcription factors including *GATA3*, *GATA4*, *GATA5*, *GATA6*, *SOX5*, *SOX6*, *TBX6*, *HAND1*, *HAND2*, and *ISL1*. From day 5 onwards, expression patterns associated with cardiac progenitor specification emerged, including activation of the cardiac transcription factors *NKX2-5*, *TBX5*, and *MEF2C*, together with the sarcomeric gene *TNNT2* (Fig. 3C).

Consistent with the WNT modulation strategy used to induce cardiac mesoderm, we observed dynamic regulation of canonical WNT signaling during early differentiation. WNT-responsive and feedback-associated genes, including *FAM53B*, *DKK1*, *NKD1*, and *AXIN2*, were upregulated, while several WNT modulators, including *SFRP1*, *TOB1*, and *TCF7L2*, showed reduced expression (Fig. 3D, Supp Fig. 2B).

In parallel, components of the FGF signaling pathway, including *FGF4*, *FGF18*, and *FGF19*, were specifically upregulated, consistent with the reported expression in the primitive streak during gastrulation^46^. FGF-associated transcriptional activity persisted during differentiation through days 3 and 5. In addition, the upregulation of Hedgehog-responsive genes, including *FOXF2* and *PTCH1*, supports activation of Hedgehog signaling during early cardiac mesoderm specification ^47,48^ (Fig. 3D, Supp Fig. 2B).

Together, these transcriptional and signaling dynamics recapitulate the progression from a shared primitive streak/mesendoderm state toward lineage-specific endodermal and mesodermal developmental trajectories, while highlighting stage-specific transcriptional programs associated with the transition from early mesendodermal competence to lineage commitment (Fig. 3E).

### Endoderm-biased genes define transcriptional, signaling, and metabolic features of definitive endoderm specification

To identify genes potentially involved in definitive endoderm specification, we filtered for genes strongly upregulated in the endodermal lineage at day 3, using a threshold of logFC ≥ 2 and adjusted P < 0.05, while excluding genes with comparable induction in the mesodermal lineage. This approach prioritized genes preferentially expressed during endodermal differentiation relative to mesodermal trajectory. Comparison of their temporal expression profiles with canonical endoderm makers showed that the candidate regulators, retinoic acid-associated gene *ALDH1A3*, and selected metabolic genes, exhibited a similar pattern of endoderm-specific induction, supporting their association with definitive endoderm specification (Fig. 4A, 4B).

**Fig 4A.**
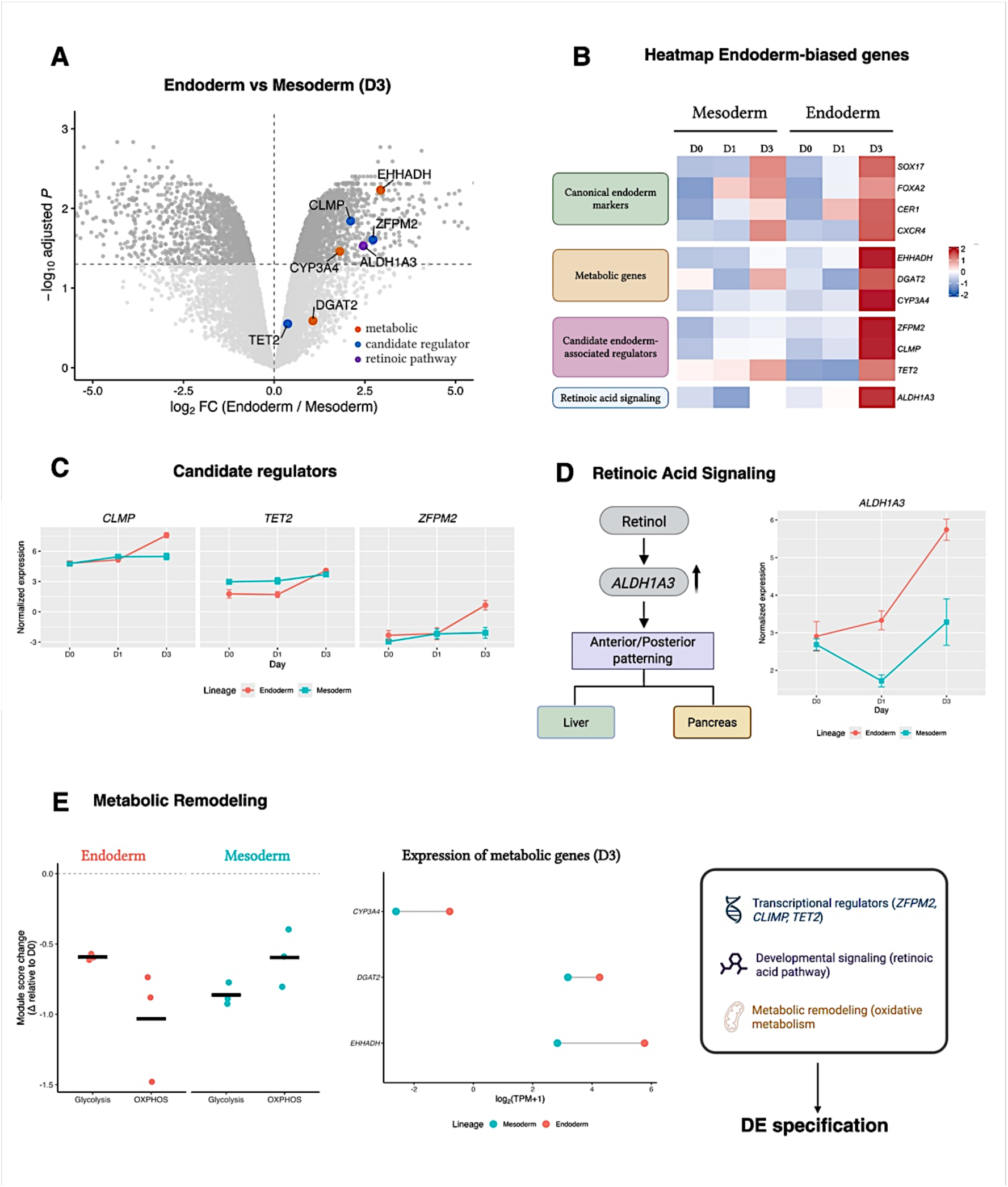
Volcano plot showing differential expression between endoderm and mesoderm lineages at day 3. Genes identified as endoderm-biased based on lineage-specific differential expression analysis (logFC ≥ 2, adjusted *P* < 0.05 in endoderm and not strongly induced in mesoderm) are highlighted. Candidate developmental regulators are shown in blue *ZFPM2, TET2, CLMP*, the retinoic acid-associated gene *ALDH1A3* is shown in purple, and metabolic genes *EHHADH, CYP3A4, DGAT2* are shown in orange. **Fig 4B | Heatmap showing normalized expression (z-score) of canonical endoderm markers, candidate endoderm-associated regulators, retinoic acid signaling, and metabolic genes across mesoderm and endoderm differentiation.** Candidate regulators and *ALDH1A3* display temporal expression patterns that closely mirror canonical endoderm markers, supporting their association with definitive endoderm specification. **Fig 4C | Temporal expression profiles of candidate endoderm-associated regulators (*ZFPM2, CLMP, TET2*) during differentiation.** Expression values represent mean normalized expression ± SEM across biological replicates. Endoderm samples are shown in **red** and mesoderm samples in **blue**. **Fig 4D | Retinoic acid signaling.** Schematic representation of retinoic acid biosynthesis and downstream developmental patterning pathways. The accompanying plot shows *ALDH1A3* expression across mesoderm and endoderm differentiation. **Fig 4E | Metabolic remodeling associated with endoderm differentiation.** Left, glycolytic and oxidative phosphorylation module score changes relative to day 0, showing distinct metabolic states in endoderm (**red**) and mesoderm (**blue**) at day 3. Center, expression of representative metabolic genes (***EHHADH, DGAT2, CYP3A4***) in mesoderm and endoderm at day 3. Right, summary model integrating candidate transcriptional regulators, retinoic acid signaling, and metabolic remodeling as coordinated features associated with definitive endoderm specification.

This analysis identified additional developmental regulators not classically associated with definitive endoderm differentiation, including *ZFPM2* (*FOG2*), *CLMP* and *TET2*. *ZFPM2* and *CLMP* displayed pronounced lineage-restricted induction during endoderm differentiation. *TET2* showed a more gradual increase in expression within the endoderm lineage, whereas expression remained relatively stable in mesoderm, indicating lineage-specific temporal regulation despite a more modest endpoint difference. (Fig. 4C). These genes may represent new candidate regulators or markers associated with definitive endoderm lineage specification.

In addition, genes associated with retinoic acid signaling were enriched in the endoderm-biased gene set, including *ALDH1A3* (Fig. 4D), which encodes an enzyme involved in retinoic acid synthesis. Given the established role of retinoic acid signaling in anterior–posterior endoderm patterning and foregut-derived organ specification, including liver and pancreas development, these findings suggest that retinoic acid-associated transcriptional programs may contribute to early endodermal patterning in our differentiation system^49–51^.

Finally, the endoderm-enriched gene set included several metabolic enzymes associated with lipid, glycogen, and energy metabolism, including *EHHADH*, *DGAT2*, and *CYP3A4* (Fig. 4E). This pattern is consistent with our previous module-score analysis, which indicated that endodermal differentiation is associated with a relative enrichment of oxidative phosphorylation-related transcriptional features compared with the mesodermal trajectory, which retained stronger glycolysis-associated signatures (Fig 4E).

Together, these findings suggest that transcriptional regulation, developmental signaling pathways, and metabolic state collectively contribute to definitive endoderm lineage establishment. This analysis identifies a set of endoderm-biased genes, including candidate regulators and pathway-associated markers, that help define the transcriptional landscape associated with endoderm lineage commitment.

## DISCUSSION

In this study, we show that comparative transcriptomics can resolve developmental intermediates that may be overlooked when differentiation is assessed solely using endpoint lineage markers. Using transcriptomic datasets from differentiation trajectories representing the three embryonic germ layers, we first established the global relationships between developmental trajectories. Because ectoderm rapidly diverges from mesendoderm-derived lineages under directed differentiation conditions, it was used as a developmental reference to define early lineage separation, while subsequent analysis focused on the shared mesendondermal program and the transcriptional mechanisms underlying mesoderm and endoderm divergence. By resolving differentiation trajectories over time, we identified distinct transcriptional programs associated with the transition from early mesendondermal competence to lineage commitment, together with candidate regulators and cellular-state-associated programs linked to mesodermal and endodermal specification.

We first focused on the initial steps of gastrulation, during which the primitive streak (PS) emerges from thickening of the epiblast^7^. Before this developmental landmark, all three germ layers originate from a common epiblast progenitor. Although the three germ-layer trajectories were generated using independent differentiation protocols, PCA analysis revealed a broadly similar temporal progression, consistent with their shared epiblast origin. Alongside this, a clear early segregation of the ectodermal trajectory from the mesendodermal trajectories was already evident. Neuroectodermal samples were clearly separated from both mesodermal and endodermal samples, consistent with the developmental origin of ectoderm from anterior primitive ectoderm cells that do not undergo epithelial-to-mesenchymal transition (EMT) and therefore do not ingress through the PS^7,52^. These ectodermal progenitors later give rise to neuroectoderm and surface ectoderm. In mouse development, this segregation occurs earlier than the complete establishment of both definitive endoderm and cardiac mesoderm ^52^. In contrast, mesodermal and endodermal trajectories remained transcriptionally close, consistent with their shared emergence through a primitive streak-associated mesendodermal intermediate and supporting previous reports of transient convergence between these two differentiation trajectories^5,31–33^.

Focusing on this early clustering showed that mesodermal and endodermal samples shared transcriptional similarities during the first days of differentiation, reminiscent of a common PS-associated origin and mesendodermal competence state. Heatmap analysis showed close proximity between D1 samples from both differentiation protocols, together with enrichment for gastrulation- and patterning-associated biological processes. This was accompanied by co-expression of previously described PS/mesendoderm markers, including *TBXT*, *MIXL1*, *EOMES*, *CER1*, and *GSC*. Mesoderm-associated markers, such as *HAND1*, *TWIST1*, *MSX2*, and *TBX3*, and endoderm-associated markers, such as *SOX17* and *GATA6*, were also co-expressed during these initial stages. These genes may therefore represent not only lineage-associated markers, but also components of a transient mesendodermal competence state preceding stable lineage commitment. In agreement with these findings, a previous study of guided mesendoderm induction in hiPSCs reported that early differentiation stages, from 0 to 48 h, retain substantial transcriptional plasticity, with many cells co-expressing primitive streak, mesodermal, and endodermal markers^34^.

Subsequently, our data showed that D3 endodermal samples retained greater transcriptional proximity to earlier differentiation stages than mesodermal samples, suggesting that although both lineages initially traverse a shared mesendodermal state, their subsequent commitment dynamics diverge. This pattern is consistent with more rapid stabilization of the endodermal transcriptional identity, whereas mesodermal differentiation remains transcriptionally dynamic during early specification. Interestingly, during embryogenesis, definitive endoderm specification occurs after anterior mesoderm progenitors delaminate from the primitive streak^4^. Thus, the transcriptional differences observed here may reflect lineage-specific cellular-state stabilization and remodeling, rather than a direct recapitulation of the exact temporal sequence of embryonic lineage emergence.

Module-scoring analysis revealed an earlier reduction in proliferation-associated programs and lower chromatin-remodeling-associated transcriptional scores in endodermal samples, whereas mesodermal samples retained signatures consistent with ongoing transcriptional remodeling. Higher chromatin-remodeling-associated scores in mesoderm may reflect the progressive establishment of cardiac progenitor identity. This interpretation is consistent with previous studies showing that cardiac mesoderm specification is accompanied by extensive chromatin remodeling that extends beyond initial lineage commitment and supports the sequential activation of cardiac regulatory networks^6,22,37^. In contrast, lower chromatin-remodeling-associated scores in endoderm may reflect earlier stabilization of lineage-specific transcriptional programs following definitive endoderm induction by NODAL/Activin A signaling. Together, these findings support the notion that mesoderm specification involves a prolonged period of developmental plasticity, whereas endoderm transitions more rapidly toward a stabilized lineage identity.

Beyond differences in transcriptional plasticity, our analyses also revealed lineage-specific metabolic signatures. Increasing evidence suggests that metabolic remodeling is closely coupled to cell-fate decisions, with changes in glycolytic and mitochondrial programs contributing to both epigenetic regulation and lineage commitment. Importantly, metabolic and chromatin-associated remodeling are increasingly recognized as interconnected processes, as mitochondrial metabolism influences the availability of metabolites required for epigenetic regulation and transcriptional control during lineage specification^21,53^. We therefore asked whether the divergent differentiation dynamics observed between mesoderm and endoderm were accompanied by distinct metabolic transcriptional states. Our data suggest that endoderm displays relatively stronger OXPHOS-associated transcriptional features, whereas mesoderm shows a less pronounced reduction in glycolysis-associated programs. Glycolysis has been shown to play a critical role during early mesendoderm specification by promoting NODAL, WNT, and FGF signaling, whereas inhibition of glycolysis favors neuroectodermal differentiation^54^. Consistent with this developmental framework, both mesodermal and endodermal trajectories displayed similar glycolysis-associated signatures during the earliest stages of differentiation, reflecting their shared primitive streak-associated origin.

However, endodermal differentiation showed relative enrichment of oxidative phosphorylation-associated transcriptional programs and metabolic genes involved in energy homeostasis. Emerging evidence indicates that metabolic remodeling contributes to definitive endoderm specification, with mitochondrial activity and oxidative metabolism supporting lineage commitment through both bioenergetic and epigenetic mechanisms. Consistent with this model, disruption of mitochondrial homeostasis impairs definitive endoderm differentiation, highlighting the importance of mitochondrial function during endoderm establishment^18,19^. Together, these observations suggest that the metabolic signatures observed in our dataset are associated with definitive endoderm commitment, rather than merely changes in cellular energetic demand. In line with evidence linking metabolism to cell-fate regulation, our findings support the view that metabolic remodeling is associated with the regulatory networks that shape developmental fate decisions.

In addition to the cellular-state transitions described above, our transcriptomic analyses recapitulated key developmental signaling pathways known to govern germ-layer specification. Endodermal differentiation was associated with activation of NODAL/TGF-β-responsive programs^40–42^, whereas mesodermal differentiation displayed dynamic regulation of WNT-, FGF-, and Hedgehog-associated transcriptional networks^45–47^. These observations are consistent with the established roles of these pathways during primitive streak formation, mesendoderm specification, and cardiac mesoderm development, further supporting the developmental relevance of the differentiation systems used in this study.

Beyond canonical endoderm regulators, our analysis identified several genes not typically associated with definitive endoderm specification, including *ZFPM2*, *CLMP*, and *TET2*. Although their roles in endoderm differentiation remain poorly defined, each has established functions in embryonic development that may provide clues to their potential involvement in lineage progression. *ZFPM2*, also known as *FOG2*, functions as a cofactor for GATA transcription factors and is required for cardiac, pulmonary, and diaphragmatic development, raising the possibility that it may modulate GATA-dependent transcriptional networks active during endoderm differentiation^55,56^. CLMP is a cell-adhesion molecule implicated in intestinal morphogenesis, with loss-of-function mutations causing congenital short bowel syndrome, suggesting a potential role in the epithelial organization programs that accompany endoderm development^57^. TET2, an epigenetic regulator involved in DNA demethylation and developmental lineage transitions, may similarly contribute to chromatin-associated regulatory events linked to the establishment of endoderm identity^58,59^.

Functional studies will be required to determine whether these genes play causal roles in endoderm specification. Nevertheless, their enrichment in our dataset highlights additional candidate pathways and regulatory mechanisms that may contribute to human endoderm differentiation and maturation. The enrichment of *ALDH1A3* and other retinoic acid-associated genes further suggests activation of patterning programs linked to foregut and hepatic development^49,51,60^. Likewise, the induction of metabolic genes such as *EHHADH*, *DGAT2*, and *AGL* is consistent with the emergence of lineage-specific metabolic programs and may reflect early acquisition of hepatic competence.

Together, our results suggest that mesodermal and endodermal differentiation initially proceed through a shared mesendodermal state before diverging through distinct cellular-state trajectories. Endodermal differentiation was characterized by earlier transcriptional stabilization, reduced chromatin-remodeling-associated signatures, and relative enrichment of oxidative metabolism-associated transcriptional programs. In contrast, mesodermal differentiation retained features consistent with prolonged developmental plasticity and progressive cardiac specification. These findings highlight that lineage commitment is shaped not only by developmental transcription factors, but also by coordinated changes in proliferation, metabolism, and chromatin-associated regulatory programs. By comparing differentiation trajectories across germ layers, our approach resolved transient intermediate states and lineage-specific cellular-state dynamics. A schematic model integrating the developmental and transcriptomic features identified in this study is presented in Figure 5 (Fig. 5). Beyond providing mechanistic insight, the markers obtained from these transient intermediate-states may represent candidate molecular checkpoints for monitoring differentiation progression. Their expression could be assessed using simple target assays, such as RT-qPCR, to confirm expected developmental trajectory during bioprocessing. Future studies should determine whether these intermediate markers can reliably predict final differentiation efficiency.

**Figure 5.**
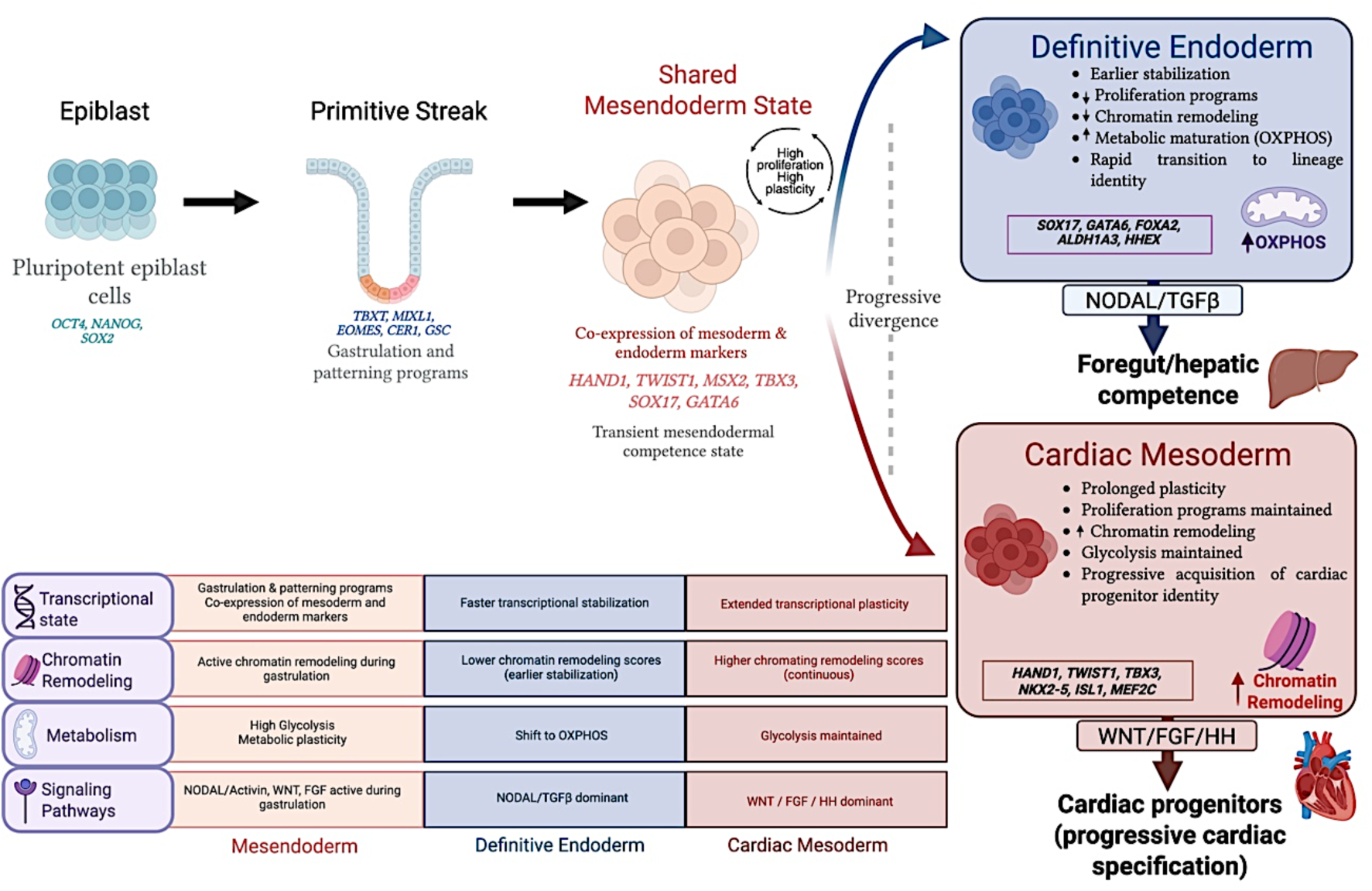
Summary model of transcriptomic divergence during mesendoderm specification. Mesodermal and endodermal differentiation initially proceed through a shared primitive streak-associated mesendodermal state before diverging into distinct developmental trajectories. Endoderm undergoes earlier transcriptional stabilization and enrichment of oxidative phosphorylation-associated programs, whereas mesoderm displays prolonged transcriptional plasticity, continued chromatin remodeling, and progressive cardiac specification. The schematic integrates the major transcriptomic signature on canonical markers, metabolic, chromatin-associated, and signaling features identified across the differentiation trajectories.

Despite these insights, several limitations should be considered. First, the analyses performed here were based on bulk RNA-seq datasets, which do not resolve cellular heterogeneity. As a result, they cannot distinguish whether the transient mesendodermal transcriptional states identified reflect individual cells co-expressing mesodermal and endodermal programs, or mixed populations of partially committed cells. Future single-cell RNA-seq analyses would help refine the temporal and cellular resolution of these states.

Second, cellular states were inferred exclusively from transcriptomic data. Although our interpretations are supported by previous literature, complementary approaches such as Assay for Transposase-Accessible Chromatin using sequencing (ATAC-seq), chromatin profiling, metabolomics, and metabolic flux analysis would further support causal relationships between these processes and lineage commitment. In addition, the 2D differentiation systems used here capture molecular aspects of early development but do not fully reproduce the effects of spatial organization, morphogen gradients, and cell-to-cell communication. Therefore, the molecular processes identified likely represent only a subset of those occurring during gastrulation and early embryogenesis. Finally, functional validation of candidate regulators through targeted perturbation studies would further determine whether these genes actively participate in lineage commitment.

## CONCLUSIONS

In this study, we applied comparative transcriptomic analysis to directed differentiation models representing derivatives of the three embryonic germ layers. Our results revealed a transient primitive streak-like mesendodermal state shared by mesodermal and endodermal trajectories prior to lineage divergence, supporting the conservation of key developmental transitions in vitro. We further show that lineage commitment is accompanied by distinct cellular-state dynamics, with endoderm displaying earlier transcriptional stabilization and mesoderm retaining features consistent with progressive specification and developmental plasticity. These differences were reflected not only in lineage-associated transcription factors, but also in coordinated proliferation-associated, chromatin-remodeling-associated, and metabolic transcriptional programs. Together, these findings demonstrate that comparative transcriptomics can resolve developmental intermediates and cellular-state transitions during human pluripotent stem cell differentiation, providing a broader framework for evaluating lineage commitment beyond canonical marker expression.

## ACKNOWLEDGEMENTS

We acknowledge funding received from Fundação para a Ciência e a Tecnologia (FCT), through Institute for Bioengineering and Biosciences (projects UIDB/04565/2020 and UIDP/04565/2020), through Associate Laboratory Institute for Health and Bioeconomy (LA/ P/0140/2020), and project Mini-Hearts (PTDC/EMD-TLM/29728/2017). FCT is also acknowledged for the PhD grants to ACB, MAB, JPC and ARG (UI/BD/153364/2022, SFRH/PD/BC128362/2017, SFRH/PD/BD135500/2018 and SFRH/PD/BD/128373/2017, respectively).

## AUTHOR CONTRIBUTIONS

All authors contributed to the conceptualization and design of the study. M.A.B., J.P.C .and A.R.G. performed the cell culture experiments. J.E.S. and L.M.M. performed the transcriptomic data acquisition and processing. A.C.B carried out the transcritpomic analyses. A.C.B. and T.G.F. interpreted the data and wrote the manuscript. M.A.B., A.R.G.,J.E.S., L.M.M, J.M.S.C., D.M. and M.M.D. provided critical feedback and contributed to revising the manuscript. All authors read and approved the final manuscript.

## SUPPLEMENTARY FIGURES

**Supp 1A.**
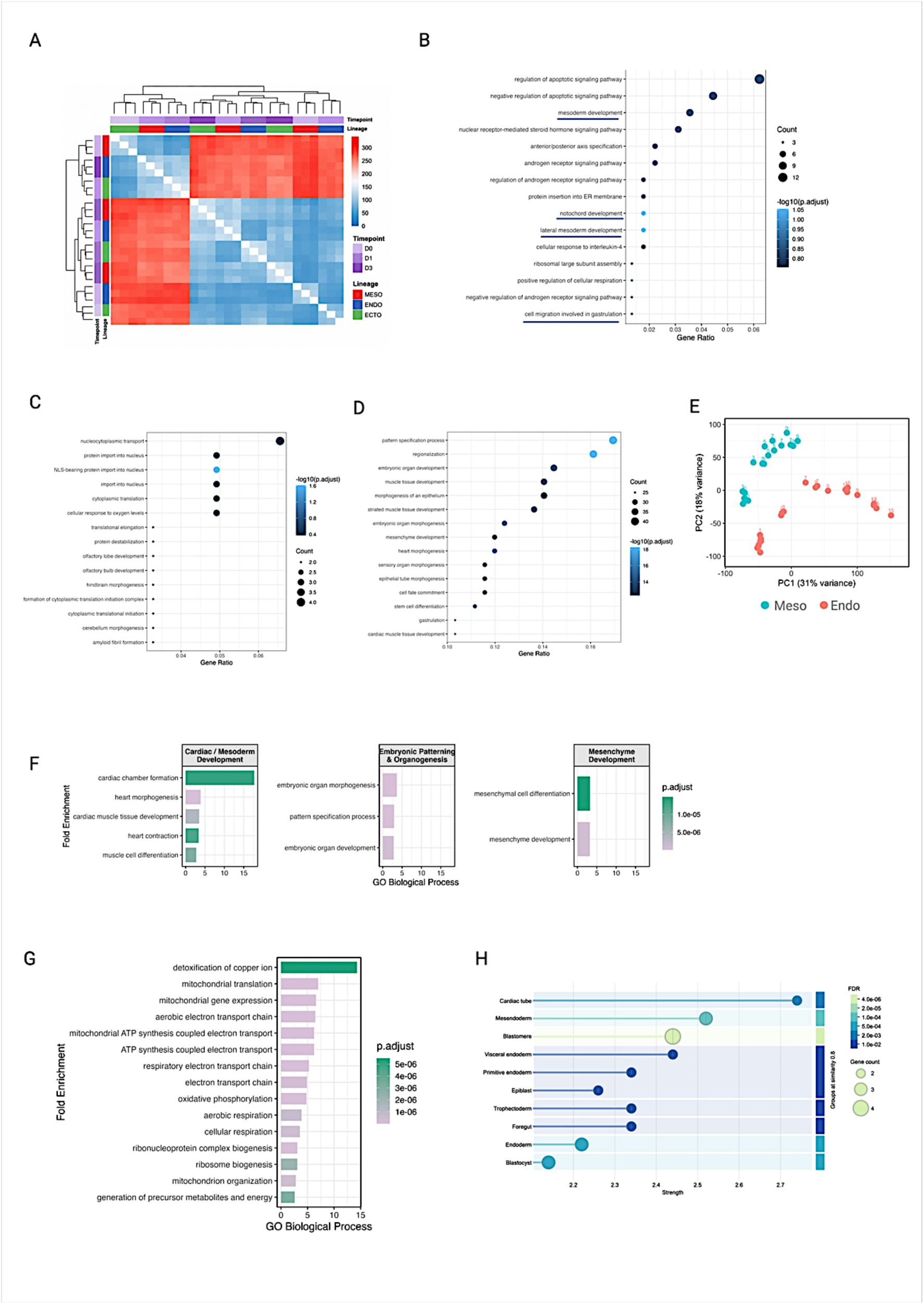
Hierarchical clustering of D0, D1, and D3 transcriptomes based on sample-to-sample Euclidean distances calculated from normalized gene expression profiles: Samples cluster primarily by lineage, with endoderm, mesoderm, and ectoderm forming distinct groups. Annotation bars indicate lineage identity and differentiation timepoint. **Supp 1B | GO biological process enrichment analysis of genes contributing positively to PC1:** Dot size indicates the number of genes associated with each term, and color denotes enrichment significance (Benjamini–Hochberg adjusted p value). Terms are ordered according to gene ratio. **Supp 1C | GO biological process enrichment analysis of genes contributing negatively to PC1:** Dot size indicates the number of genes associated with each term, and color denotes enrichment significance (Benjamini–Hochberg adjusted p value). Terms are ordered according to gene ratio. **Supp 1D | GO biological process enrichment analysis of genes contributing negatively to PC2:** Dot size indicates the number of genes associated with each term, and color denotes enrichment significance (Benjamini–Hochberg adjusted p value). Terms are ordered according to gene ratio. **Supp 1E | PCA of hiPSC mesoderm and endoderm differentiation:** Samples are colored by lineage (mesoderm, blue; endoderm, red) and labeled by differentiation day. PC1 and PC2 explain 31% and 18% of variance, respectively. **Supp 1F | GO enrichment of PC2-upregulated genes highlights mesodermal and cardiac developmental programs:** Bars show fold enrichment and are colored by adjusted P value. **Supp 1G | GO enrichment of PC2-downregulated genes highlights oxidative phosphorylation, translation, ribosomal, and mitochondrial programs:** Bars show fold enrichment and are colored by adjusted P value. **Supp 1H | Tissue enrichment analysis of lineage-associated genes:** Tissue expression enrichment was performed using STRING on the curated gene set shown in the heatmap. Bars indicate enriched tissues, colored by false discovery rate (FDR), with circle size representing the number of contributing genes. The analysis highlights preferential expression in mesoderm- and endoderm-related tissues, consistent with early developmental lineage specification.

**Supplementary Figure 2.**
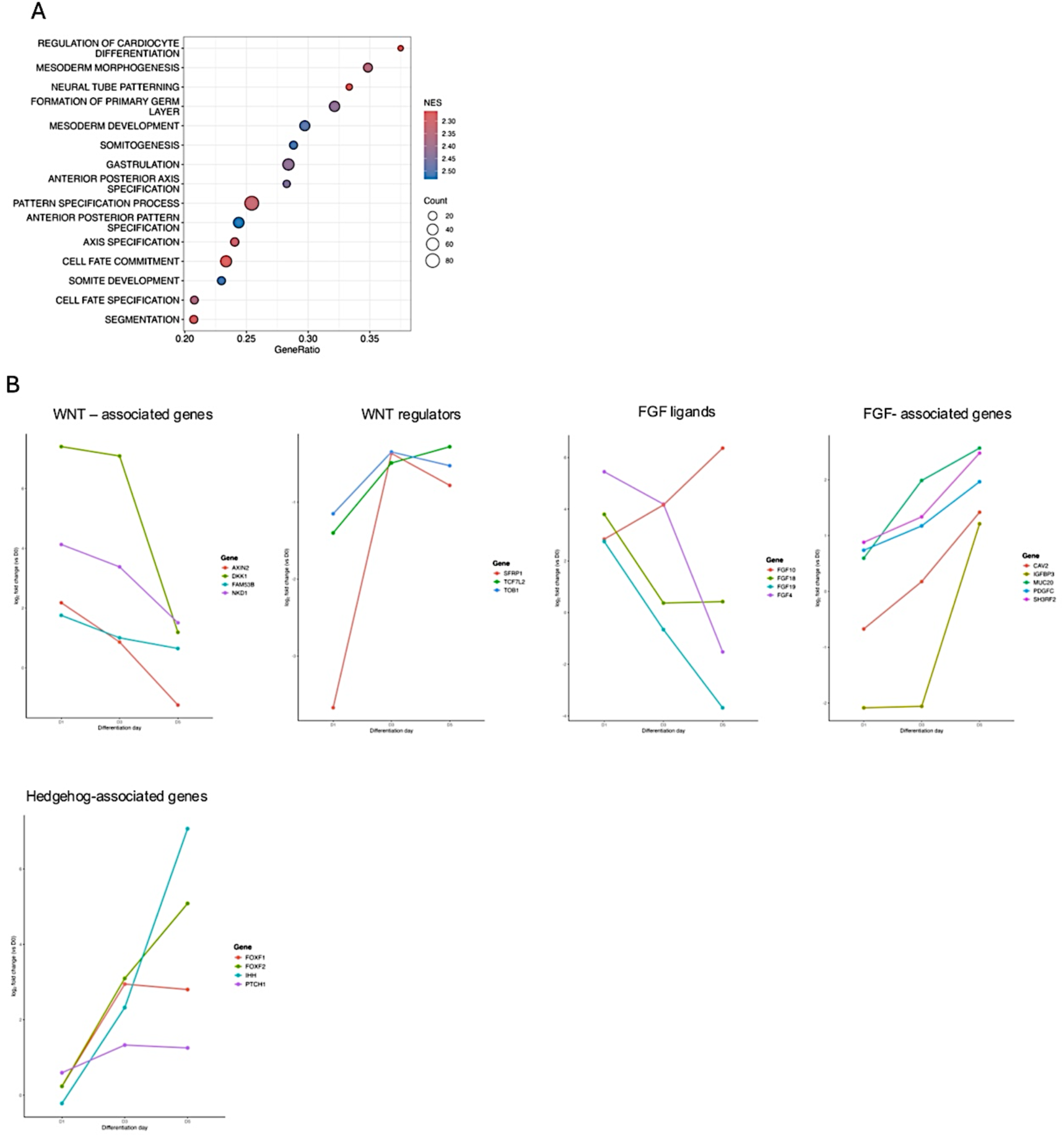
Developmental signaling pathways associated with endoderm differentiation. **(A) Gene Ontology enrichment analysis of endoderm-enriched genes highlighting developmental and cell fate specification processes.** Dot size indicates gene count and color denotes enrichment significance. **(B) Expression dynamics of representative WNT-, FGF-, and Hedgehog-associated genes across endoderm differentiation.** Expression values are shown as normalized transcript abundance over time.

